# Unitization modulates recognition of within-domain and cross-domain associations: Evidence from event related potentials

**DOI:** 10.1101/465336

**Authors:** Bingcan Li, Meng Han, Chunyan Guo, Roni Tibon

## Abstract

Although it is often assumed that memory of episodic associations requires recollection, it has been suggested that when stimuli are experienced as a unit, familiarity processes might contribute to their subsequent associative recognition. We investigated the effects of associative relations and perceptual domain during episodic encoding on retrieval of associative information. During study, participants encoded compound and non-compound words-pairs, presented either to the same sensory modality (visual presentation) or to different sensory modalities (audio-visual presentation). At test, they discriminated between old, rearranged, and new pairs while event related potentials (ERPs) were recorded. In an early ERP component, generally associated with familiarity processes, differences related to associative memory only emerged for compounds, regardless their encoding modality. In contrast, in a later ERP component associated with recollection, differences related to associative memory emerged in all encoding conditions. These findings may indicate that episodic retrieval of compound words can be supported by familiarity-related processes, regardless of whether both words were presented to the same or different sensory modalities.

## 1 Introduction

Remembering episodic associations—that several objects, people, or actions were experienced conjointly—is a core cognitive function. Such associations can involve a single sensory modality (the looks of the two people you’ve just met) or multiple sensory modalities (the sight and taste of a sandwich you’ve just ate); they can involve items that were already experienced together before (two of your friends from grad-school sitting together), or items that are now experienced together for the first time (your friend from grad-school walking down the street with your mother). Unitization—the creation of a new unit from individual items—has been proposed as an effective strategy for remembering episodic associations (e.g., Gobet et al., 2001; Graf & Schacter, 1989; Yonelinas, 1997; Yonelinas, Kroll, Dobbins, & Soltani, 1999), but the conditions that enable unitization require further specification. In the current study, we used EEG to examine the effect of different encoding conditions on unitization and on the time course of associative retrieval.

Memory for episodic associations can be assessed with an associative recognition task: following a study phase, in which stimulus-pairs are learned, participants are presented with two items, and are asked to discriminate between studied pairs (old) and studied items in new combinations (rearranged). As the individual items of old and rearranged pairs are equally familiar, differences between these conditions are attributed to associative (rather than item) memory. The widely supported dual-process theory of episodic memory posits that recognition tasks involve two separable processes: familiarity and recollection. Familiarity is a feeling of having encountered something or someone before, without retrieval of additional information, whereas recollection further provides contextual details about that encounter. This distinction is supported by evidence from many behavioral, neurophysiological, and neuroimaging studies, including event-related potential (ERP) studies showing that two qualitatively distinct ERP components are associated with recognition judgments. The first is the early mid-frontal component, showing greater negative deflection for new vs. old items, arising 300-500 ms post-stimulus presentation. This effect has been widely described as the putative electrophysiological correlate of familiarity. The second is the late posterior component, showing greater positive deflection for old vs. new items, and is prominent over left parietal electrodes 500-800 ms post-stimulus presentation. This effect is considered to be an electrophysiological correlate of recollection (reviewed by Mecklinger, 2000, Rugg & Curran, 2007, Wilding & Ranganath, 2011).

While it is generally agreed that recognition of single items can be supported by both recollection and familiarity, it has traditionally been asserted that recollection is required to retrieve episodic associations, and that associative recognition is not accessible via familiarity processes (e.g., Donaldson & Rugg, 1998, Hockley & Consoli, 1999, Yonelinas, 1997). In recent years, however, research has suggested that familiarity can contribute to associative recognition when pairs of items are unitized (i.e., treated as a single unit rather than as a pairing of two separate items). This notion is supported by a growing body of evidence (for review, see Mecklinger & Jäger, 2009; Yonelinas, Aly, Wang, & Koen, 2010), including behavioral studies (e.g., Ahmad & Hockley, 2017; Diana, Yonelinas, & Ranganath, 2008; Parks & Yonelinas, 2015; Robey & Riggins, 2017; Shao, Opitz, Yang, & Weng, 2015; Tibon, Greve, & Henson, 2018; Tibon, Vakil, Goldstein, & Levy, 2012; Tu, Alty, & Diana, 2017) and functional magnetic resonance imaging (fMRI) studies (Bader, Opitz, Reith, & Mecklinger, 2014; Diana et al., 2009; Ford, Verfaellie, & Giovanello, 2013; Haskins, Yonelinas, Quamme, & Ranganath, 2008; Memel & Ryan, 2017), as well as electrophysiological studies showing modulation of the mid-frontal ERP effect associated with familiarity, following unitization encoding (e.g., Bader, Mecklinger, Hoppstädter, & Meyer, 2010; Diana, Van den Boom, Yonelinas, & Ranganath, 2011; Guillaume & Etienne, 2015; Jäger, Mecklinger, & Kipp, 2006; Jäger, Mecklinger & Kliegel, 2010; Kamp, Bader, & Mecklinger, 2016; Rhodes & Donaldson, 2008; Tibon, Ben-Zvi, & Levy, 2014; Tibon, Gronau, Scheuplein, Mecklinger, & Levy, 2014; Zheng, Li, Xiao, Broster, & Jiang, 2015).

Two major theoretical frameworks account for mnemonic unitization effects. The first is the Domain Dichotomy (DD) view (Mayes et al., 2007). This view extends earlier neurocognitive models of recognition memory, which associate the hippocampus with recollection and the perirhinal cortex with familiarity (for review see Yonelinas, 2002), and proposes that the perirhinal cortex can mediate familiarity-based retrieval of intra-item and within-domain associations (e.g. face–face associations), but that the hippocampus mediates retrieval of cross-domain associations (e.g. scene–sound, face–voice etc.), which require recollection. In the context of unitization, this distinction implies that familiarity of unitized associations is limited to within-domain items. The second theoretical framework is the Levels of Unitization (LOU) account, which refers to the idea that there is a continuum along which associations can be unitized (Parks & Yonelinas, 2015). According to LOU, unitization is critically determined by the way in which individuals process the incoming stimuli (e.g., as a compound word or as two separate words). As such, it is assumed to operate at a fairly abstract level. Therefore, LOU predicts that it should be possible to unitize across different stimulus modalities or domains, such as words and faces or visual and auditory stimuli.

The discrepancy in the predictions posed by the two theoretical frameworks is also evident in empirical results. Indeed, while some studies support the DD view, by showing that familiarity / non-hippocampal based retrieval is useful in supporting retrieval of within-domain, but not cross-domain associations (Bastin, Van der Linden, Schnakers, Montaldi, & Mayes, 2010; Borders, Aly, Parks, & Yonelinas, 2017; Mayes et al., 2004; Tibon et al., 2014; Troyer, D’Souza, Vandermorris, & Murphy, 2011; Vargha-Khadem, Gadian, Watkins, Connelly, van Paesschen, & Mishkin, 1997), others agree with the LOU account by showing no differences between within- and cross-domain associations in familiarity based retrieval (Park & Rugg, 2011; Turriziani, Fadda, Caltagirone, & Carlesimo, 2004), or even a reversed pattern (i.e., greater familiarity based retrieval of cross-domain associations; Harlow, Mackenzie, & Donaldson, 2010; Parks & Yonelinas, 2015).

Parks and Yonelinas (2015) speculate that the complexity of the stimuli might be an important factor in determining why some but not other within-domain stimuli would benefit from unitization, and suggest that complex stimuli may be more difficult to unitize within domain because they impose greater attentional processing demands and/or because their combination does not result in a coherent unit. We propose that another possible source of this discrepancy is the nature of the associative relations of the to-be-unitized stimuli. As was mentioned above, the LOU account suggests that stimulus-pairs can vary along a continuum of their associative relations. On one end of this continuum are stimuli that already share strong associative relationships prior to the experiment, such as word compounds (e.g., “cottage pie” or “traffic jam”) or related object-pictures presented in a coherent spatial configuration (e.g., a lamp over a desk). On the other end, are stimuli that do not share any obvious relations prior to the experiment, such as semantically unrelated word-pairs or object-pictures. Crucially, pre-experimental associative relations often apply to items that are experienced via the same sensory modality (e.g., in most cases, both words comprising a compound would be presented in the same manner). Therefore, we propose that when stimuli that hold pre-existing relations are encoded within-domain, their relations are easily processed, possibly allowing within-domain (but not cross-domain) stimulus-pairs to benefit from unitization.

Accordingly, the current study examined the effects of different encoding modes, i.e., perceptual domain (within-domain, cross-domain) and word-pair type (compound, non-compound), on retrieval processes supporting associative episodic recognition, as indexed by differences in ERP correlates of retrieval. Memoranda were either compound or non-compound word pairs, presented either visually or in an audio-visual presentation. At study, participants performed a relatedness judgment for each word-pair. At test, they discriminated old, rearranged and new pairs while EEG was recorded, enabling us to examine the time courses of episodic retrieval. We set our experiment to test three main predictions. First, because compound words share strong associative relationships, and are thus located at the higher end of the unitization continuum (according to the LOU account), we predicted greater unitization effects for compounds than non-compounds. Second, with respect to unitization effects within/across perceptual domains, we predicted that if the DD view is correct, then greater unitization effects will emerge for within-domain pairs, but that if the LOU account is correct, unitization effects will not differ for within-domain and cross-domain pairs. Third, given our proposal that pre-existing relations are easily processed, allowing within-domain but not cross-domain stimulus-pairs to benefit from unitization, we predicted greater unitization effects for compounds vs. non-compounds for within-domain than for cross-domain pairs.

## 2 Methods

### 2.1 Participants

Twenty right-handed students from Capital Normal University were paid ¥30 per hour to take part in the study. Data from three participants were excluded due to insufficient artifact-free trials in one or more conditions (n < 18) following EEG artifacts removal. Seventeen participants remained (11 women, *mean age* = 23.5, range = 20 – 26 years). All participants were native Chinese speakers, had normal or corrected to normal vision, and provided informed consent as approved by The Human Research Ethics Committee of Capital Normal University.

### 2.2 Stimuli

Study stimuli included a list of 600 Chinese 2-character-word-pairs [mean total number of strokes: 8.72 (range, 4–23), mean word frequency: 98.4 (range, 3.5–493.2) occurrences per million (Liu, 1990)], divided into two sub-lists. The first sub-list included 300 compounds (e.g., ‘American movie’) and the second sub-list included 300 non-compound word-pairs with no associative or semantic relationship.

To verify word assignment into sub-lists, we conducted a pilot study based on a protocol used in previous studies (Kriukova, Bridger, & Mecklinger, 2013; Rhodes & Donaldson, 2007; Zheng et al., 2015), in which 10 native Chinese speakers (3 men and 7 women), who did not take part in the main experiment, were asked to judge how well the two words could be bound into a single concept using a scale from 1 (lowest ratings) to 7 (highest ratings). The results confirmed our initial assignment of word-pairs into sub-lists, and further showed that the set of compound word-pairs received higher rating (*Mean* = 5.94, *S.D*. = 0.86) than the set of non-compound word-pairs (*Mean* = 1.33, *S.D*. = 0.20), *t*(9) =16.61, *p* < 0.0001).

All word-pairs were randomly assigned into ten study-test blocks, including five within-domain blocks (visual-visual) and five cross-domain blocks (visual-auditory) (see Fig 1). Block order was interleaved, and counterbalanced across participants (such that for half the participants the first block was a within-domain block, and for the other half the first block was cross-domain block). Word-pairs were matched across conditions, with equivalent number of strokes and word frequency. Auditory words were edited using Cooledit software (*mean length* = 1077 ms, *S.D*. = 185.7 ms).

In each block, the study phase consisted of 20 compounds and 20 non-compounds. In *within-domain* blocks, both words were presented visually. In the *cross-domain* blocks, one word was presented visually, and the other was presented auditorily. The test phase of each block included 20 old pairs (presented at study), 20 rearranged pairs (constructed from words which were presented with other words at study), and 20 new word-pairs. Importantly, old and rearranged words were presented at test with their perceptual domain (within-domain, cross-domain) unchanged from study. Additionally, rearranged pairs were presented at test with their type (compound, non-compound) unchanged from study, such that compounds were rearranged into new compounds and non-compounds were rearranged into new non-compounds. For example, the compound (e.g., ‘美国-电影’ meaning ‘American-movie’) was rearranged into another compound (e.g., ‘美国-总统’ meaning ‘American-president) and the non-compound (e.g., ‘电脑-肥皂’ meaning ‘Computer-soap) was rearranged into non-compound (e.g., ‘电脑-空气’ meaning ‘Computer-air). Overall, the design included 12 conditions: 2 perceptual domains (within-domain, cross-domain) X 2 word-pair types (compound, non-compound) X 3 response types (old, rearranged, new), with 50 trials in each condition.

**Figure 1.**
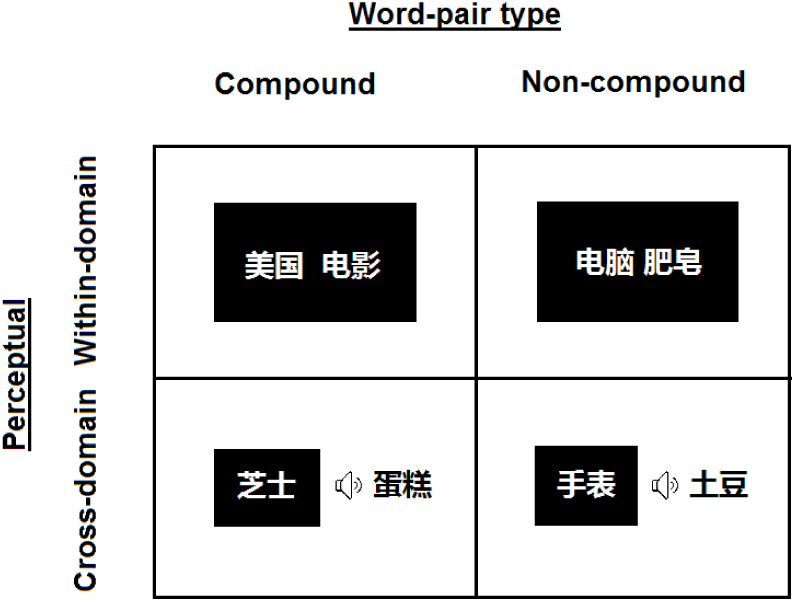
Example stimuli. Word-pair type was either ‘compound’ or ‘non-compound’. Perceptual domain was either visual for both words (within-domain) or audio-visual (with one word presented visually, and the other auditorily; cross-domain). The examples of word-pairs mean ‘American movie’ (top-left), ‘cheese cake’ (bottom-left), ‘computer soap’ (top-right) and ‘watch potato’ (bottom-right).

### 2.3 Procedure

Participants were seated in an electrically shielded and quiet room. Stimuli were displayed by Presentation software (Neurobehavioral Systems, San Francisco, CA) on a 17-inch Dell computer monitor. Visual stimuli were displayed in white 18-point Simhei font against a black background. At a viewing distance of 70 cm, the words subtended a maximum horizontal visual angle of 3.7°, and a maximum vertical visual angle of 1.4°. Auditory words were read loudly and clearly by a male radio announcer, and displayed via earphones.

During the study phase, each trial began with the presentation of a fixation cross (+) at the center of the screen for 1,000 ms, after which the word-pair was displayed. In *within-domain* blocks, both words in the word-pair were presented on the screen for 1000 ms followed by a blank screen for 500 ms. In *cross-domain* blocks, one word was presented on the screen for 1000 ms, with the auditory word concurrently presented through earphones. Participants were instructed to indicate whether the two words are related or not, by pressing marked keyboard keys using their left / right index finger. Key assignment was counterbalanced across participants.

During the test phase, each trial began with the presentation of a fixation cross for 1,000 ms. In *within-domain* blocks, both words in the word-pair were presented on the screen for 2000 ms. In *cross-domain* blocks, one word was presented on the screen for 2000 ms, and the other was concurrently presented through earphones. During the test phase, participants were asked to judge whether the same word-pair was presented during the study phase (“old”), whether the words comprising the pair were presented at study, but with different pairing (“rearranged”), or whether the words comprising the word-pair were not seen before (“new”). Participants provided their responses using the F key (with their left index finger), and the J and K keys (with their right index and middle finger). Hand assignment was counterbalanced across participants.

### 2.4 EEG data collection and preprocessing

EEG signals were recorded from 62 Ag/AgCl electrodes embedded in an elastic cap equipped with a NeuroScan SynAmps system, and preprocessed with Neuroscan software. EEG data were collected at a sampling rate of 500 Hz with a 0.05–100 Hz band pass filter. Horizontal electrooculogram (HEOG) were recorded bipolarly from electrodes placed 1 cm to the left and right of the outer canthi. Vertical electrooculogram (VEOG) were recorded bipolarly from electrodes placed above and below the left eye. All voltages were referenced to the left mastoid online and re-referenced offline to the average of the left and right mastoid. Impedance was less than 5 kΩ and EEG/electrooculogram (EOG) signals were digitally band pass filtered from 0.05 Hz to 40 Hz. The EEG was segmented into 1200 ms epochs starting from 200 ms before the presentation of the stimuli, and then subjected to baseline correction with the 200 ms window preceding the stimuli. Trials with voltage exceeding ±75 *μ* V at any electrode and other EEG artifacts other than blinks were rejected. Blink artifacts were corrected using a linear regression estimate (Semlitsch, Anderer, Schuster, & Presslich, 1986).

The EEG analyses only included trials with correct responses. Mean numbers of within-domain analyzed trials were 45.4 (old), 32.1 (rearranged), and 40.1 (new) for compounds, and 36.4 (old), 34.9 (rearranged), and 39.8 (new) for non-compounds. Mean numbers of cross-domain analyzed trials were 44.6 (old), 33.9 (rearranged), and 38.8 (new) for compounds, and 33.9 (old), 36.1 (rearranged), and 39.9 (new) for non-compounds. The minimal trial number in each condition was 20 (after excluding 3 participants who had less than 18 trials in one or more conditions, see section 2.1 above).

### 2.5 Data analyses

For inferential statistics, repeated measures analyses of variance (ANOVAs) were conducted. Subsidiary analyses were performed using repeated measures ANOVAs or t-tests as appropriate. Probability values (p-values) for follow-up analyses were adjusted by applying a Bonferroni correction. The significance level was set to *α*=.05. Given our interest in unitization effects on associative recognition (i.e., the difference between old and rearranged responses), we did not analyze new responses (although data associated with these responses is shown for completeness). Furthermore, because the current study focuses on mnemonic effect, only main effects and interactions involving the factor of response type are reported.

#### 2.5.1 Behavioral analyses

To analyze data from the study phase, repeated measures ANOVA was performed with factors of perceptual domain (within-domain, cross domain) and word-pair type (compound, non-compound) on relatedness rating (% ‘related’ responses).

To examine memory performance during test, repeated measures ANOVAs with factors of perceptual domain (within-domain, cross-domain), word-pair type (compounds, non-compound), and response type (old, rearranged) were conducted on accuracy (% correct responses) and response times (RTs). To further analyze associative discrimination, associative Pr indices (the proportion old pairs correctly classified as ‘old’, minus the proportion of rearranged pairs incorrectly classified as ‘old’) were submitted to a repeated measures ANOVA, with perceptual domain and word-pair type as within-subject factors.

#### 2.5.2 ERP analyses

Our ERP analyses includes two sections. The critical hypothesis underlying the current study was that unitized, but not un-unitized associations, may promote the contribution of familiarity to associative recognition. Therefore, in the first section, we focused on the early frontal old/rearranged effect, conventionally interpreted as the putative ERP correlate of associative familiarity, to test the three predefined predictions described in the introduction: (1) that the old/rearranged effect will be greater for compounds than non-compounds, (2) that the old/rearranged effect will be greater for within-domain than cross-domain word-pairs if the DD view is correct, but will not differ between perceptual domains if the LOU account is correct, and (3) that the difference between compounds and non-compounds in the magnitude of the old/rearranged effect, will be greater for within-domain than for cross-domain pairs.

All three predictions were tested using a single repeated measures ANOVA, with perceptual domain (within-domain, cross-domain), word-pair type (compound, non-compound) and response type (old, rearranged) as repeated factors, and with averaged data in the mid-frontal site (Fz) in an early time window (300-500), where familiarity effects are typically observed (reviewed by Friedman & Johnson, 2000; Mecklinger, 2000; Rugg & Curran, 2007; Wilding & Ranganath, 2011), as the dependent measure. For our first prediction, we expected a 2-way interaction between word-pair type and response type, such that the difference between old and rearranged responses (the old/rearranged effect) would be greater for compounds than for non-compounds. For our second prediction, we expected a 2-way interaction between perceptual domain and response type, such that the old/rearranged effect would be greater for within-domain than for cross-domain pairs, if the DD view is correct, but not if the LOU account is correct. For our third prediction, we expected a 3-way interaction between all three factors, such that the difference between compounds and non-compounds in the magnitude of the old/rearranged effect will be greater for within-domain than for between-domain word-pairs. Null results were followed-up with Bayesian analyses, performed using R version 3.4.1 as implemented by RStudio version 1.0153, with a BayesFactor package (Morey & Rouder, 2015).

In the second section of the ERP analyses, we provide comprehensive analyses of the data, which were not guided by specific predictions. For this purpose, two standard time windows were selected, 300–500 ms (early) and 500–800 ms (late), to capture the early frontal and late posterior effects, respectively. Mean amplitudes for each condition were obtained from three frontal (F3, Fz, F4), three central (C3, Cz, C4), and three parietal (P3, Pz, P4) electrodes (see Figure 2, panel E, for location map of analyzed electrodes). To investigate associative recognition effects, for each time window we ran a repeated measures ANOVA on this mean amplitude, with perceptual domain (within-domain, cross-domain), word-pair type (compound, non-compound), response type (old, rearranged), anteriority (anterior, central, posterior) and laterality (left, mid, right) as within-subject factors. Significant main effects and interactions were followed-up with post-hoc ANOVAs or paired t-tests.

**Figure 2.**
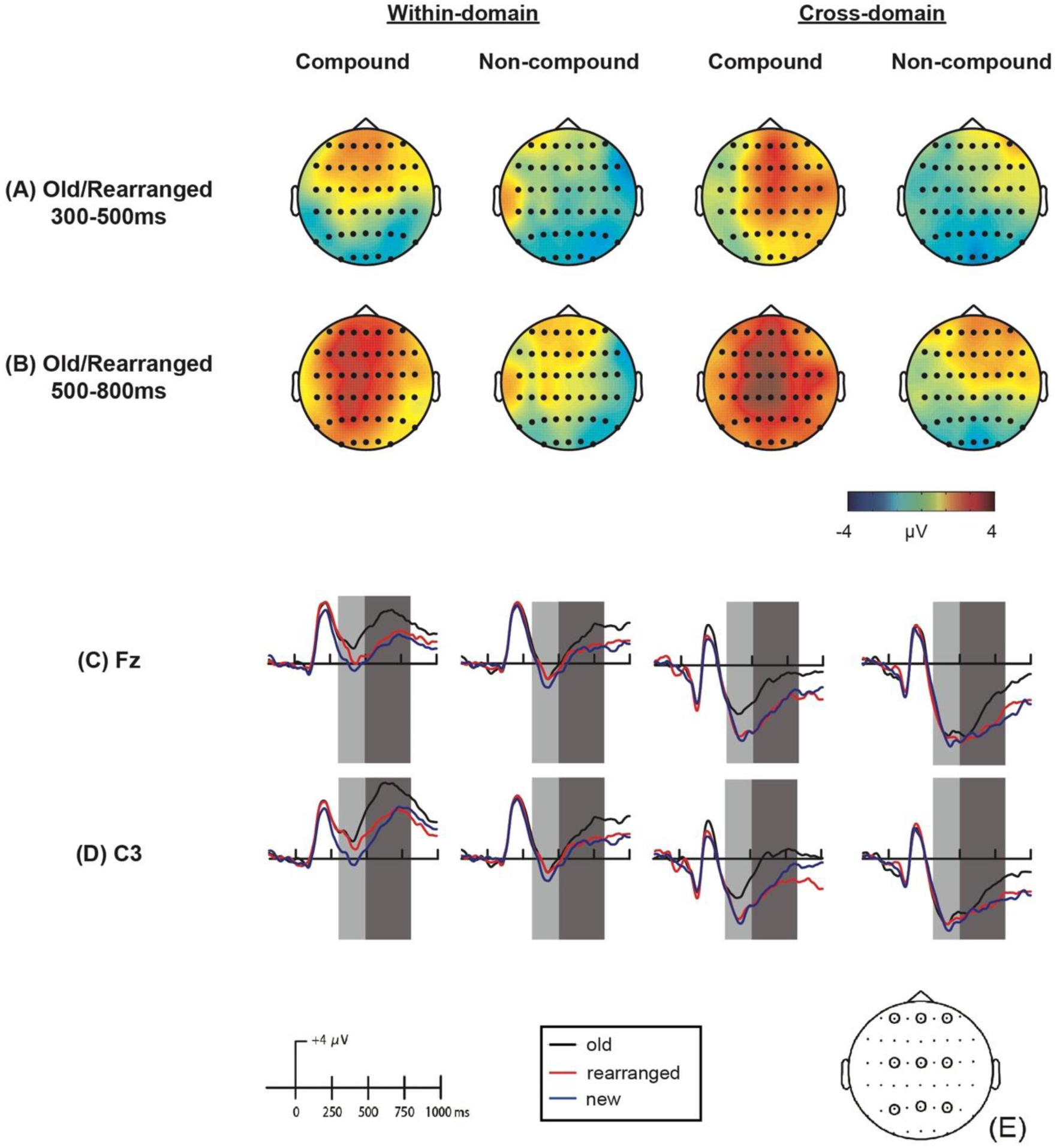
Panels A-B: topographic maps depicting the distribution of the old/rearranged effect in the early (300-500 ms; panel A) and late (500-800 ms; panel B) time windows, in the various perceptual domains (within-domain, cross-domain) and word-pair types (compound, non-compound). Panels C-D: ERP waveform for ‘old’ (black), ‘rearranged’ (red) and ‘new’ (blue; not analyzed) responses in the various perceptual domains and word-pair types at Fz (panel C) and C3 (panel D). Analyzed time windows are highlighted in light gray (300–500ms; early time window) and dark gray (500-800ms; late time window. (E) Schematic depiction of EEG channels implicated in the analyses: left frontal (F3); mid frontal (Fz); right frontal (F4); left central (C3); mid central (Cz); right central (C4); left posterior (P3); mid posterior (Pz); right posterior (P4).

## 3 Results

### 3.1 Behavioral results

#### 3.1.1 Study phase

Data from the study phase are shown in Table 1. The analysis of this data revealed a main effect of word-pair type, *F*(1, 16) = 2215.23, *p* < .0001, 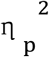 = 0.99, with increased proportion of ‘related’ responses for compound than non-compound word-pairs, and a main effect of perceptual domain, *F*(1, 16) = 28.275, *p* < .0001, 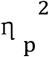 = 0.64, with increased proportion of ‘related’ responses for within-domain than cross-domain word-pairs.

**Table 1.** %‘related’ responses in the various study conditions

| Within-domain |  | Cross-domain |  |
| --- | --- | --- | --- |
| Compound | Non-compound | Compound | Non-compound |
| 97.41 (4.19) | 10.59 (10.5) | 95.65 (3.06) | 3.88 (4.65) |

Note: Standard deviations are shown in parenthesis.

#### 3.1.2 Test phase: Accuracy

Table 2 shows accuracy data at test. The analysis of % accuracy revealed a main effect of word-pair type, *F*(1, 16) = 12.35, *p* = .003, 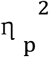 = 0.44, with better memory for compounds compared to non-compounds, and a main effect of response type, *F*(1, 16) = 90.96, *p* < .0001, 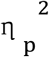 = 0.85, with better memory for old relative to rearranged word-pairs (*ps* < 0.001). Furthermore, an interaction between perceptual domain and response type was revealed, *F*(1, 16) = 6.60, *p* = .021, 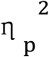 = 0.29, stemming from greater accuracy for cross-domain compared to within-domain word-pairs for rearranged responses, *t*(16) = -2.74, *p* = .015, but not for old responses, *t*(16) = -0.9, *p* = .384. In addition, an interaction between conceptual mode and response type was revealed, *F*(1, 16) = 33.13, *p* < .0001, 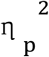 = 0.67, stemming from greater accuracy for compounds compared to non-compounds for old responses, *t*(16) = 5.9, *p* < .0001, but marginally decreased accuracy for compounds compared to non-compounds for rearranged response, *t*(16) = -1.95, *p* = 0.069.

**Table 2.** Mean accuracy (%) and associative Pr indices in the various test conditions

|  | Within-domain |  | Cross-domain |  |
| --- | --- | --- | --- | --- |
|  | Compound | Non-compound | Compound | Non-compound |
| **Old** | 95.61 (3.4) | 75.76 (15.3) | 95.53 (4.8) | 73.53 (17.1) |
| **Rearranged** | 67.47 (13.4) | 72.85 (14.4) | 72.82 (11.6) | 78.47 (13.3) |
| **New** | 84.0 (9.0) | 82.91 (10.3) | 83.65 (7.0) | 86.24 (10.0) |
| **Associative Pr** | 63.61 (15.4) | 48.62 (27.4) | 68.35 (13.9) | 52.00 (28.1) |

Note: Standard deviations are shown in parenthesis. “New” responses are shown for completeness, but were not included in the analysis

For associative Pr’s (Table 2, bottom row) the analysis revealed a main effect for word-pair type, *F*(1, 16) = 12.35, *p* = .003, 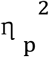 = 0.44, with greater Pr values for compounds than for non-compounds, and a marginal main effect of perceptual domain, *F*(1, 16) = 3.97, *p* = .06, 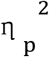 = 0.2, with greater Pr values for cross-domain than for within-domain word-pairs.

#### 3.1.3 Test phase: Reaction times

Table 3 shows mean RTs at test. The analysis for RTs revealed a main effect of perceptual domain, *F*(1, 16) = 10.09, *p* = .006,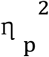 = 0.39, with faster RTs for within domain word-pairs compared to cross-domain word-pairs, a main effect of word-pair type, *F*(1, 16) = 136.81, *p* < .0001, 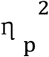 = 0.90, with faster RTs for compounds compared to non-compounds, and a main effect of response type, *F*(1, 16) = 177.83, *p* < .0001, 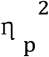 = 0.92, with faster RTs for old relative to rearranged word pairs. Additionally, an interaction between word-pair type and response type was revealed, *F*(1, 16) = 74.19, *p* < .0001, 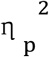 = 0.82, stemming from faster RTs for compounds compared to non-compounds for old responses, *t*(16) = -13.20, *p* < .0001, but not for rearranged response, *t*(16) = -1.47, *p* = .162.

**Table 3.** Reaction times in the various test conditions

|  | Within-domain |  | Cross-domain |  |
| --- | --- | --- | --- | --- |
|  | Compound | Non-compound | Compound | Non-compound |
| **Old** | 1173.3 (171) | 1695.2 (286) | 1264.2 (152) | 1827.7 (296) |
| **Rearranged** | 2020.7 (351) | 2055.2 (362) | 2084.2 (330) | 2152.4 (327) |
| **New** | 1557.9 (261) | 1751.8 (280) | 1722.2 (266) | 1832.5 (311) |

Note: Standard deviations are shown in parenthesis. “New” responses are shown for completeness, but were not included in the analysis

### 3.2 ERP results

#### 3.2.1 Predefined predictions

The ANOVA which was set to test our predefined predictions, revealed a significant main effect of perceptual domain, *F*(1, 16) = 104.89, *p* < .0001, 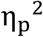= 0.87, word-pair type, *F*(1, 16) = 22.16, *p* < .0001, 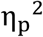= 0.58, and response type, *F*(1, 16) = 14.32, *p* = .002, 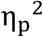= .47. Importantly, in accord with our first prediction, the analysis further revealed a significant interaction between word-pair type and response type, *F*(1, 16) = 5.05, *p* = .04, 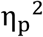= 0.24, stemming, as predicted, from a greater old/rearranged effect for compounds than non-compounds. However, the interaction between perceptual mode and response type, predicted by the DD view (but not by the LOU account), was not significant, *F*(1, 16) = 0.53, *p* = .48, 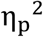= 0.03. In addition, the 3-way interaction between these factors, specified by our third prediction, was not significant either, *F*(1, 16) = 0.69, *p* = .7, 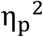= 0.01.

Thus, the ANOVA analysis yielded null results for our second and third predictions. However, as with any form of classical null-hypothesis testing, absence of evidence is not evidence of absence. We therefore used a Bayesian approach to compare null and alternate hypotheses. To test the second prediction, we collapsed across word-pair type for each participant, and subtracted the amplitude of rearranged responses from that of old responses, to calculate the magnitude of the old/rearranged effect separately for each perceptual domain. Then, we used a one-sided Bayesian t-test with a Cauchy prior scaled at sqrt(2)/2 (medium scaling), to compare two hypotheses: the LOU suggestion, that the old/rearranged effect does not differ for within-domain and cross-domain word-pairs (that is, that the standardized effect size is 0; the so-called “null hypothesis”), and the DD suggestion, that the old/rearranged effect is greater for within-domain than cross-domain word-pairs (that is, that the standardized effect size is bigger than 0). This analysis supported the null hypothesis, which was preferred by a Bayes factor of 6.35. The data thus provide moderate evidence in support of the prediction of the LOU account.

To test our third prediction, we calculated the old/rearranged effect for each participant in each condition, and subtracted the magnitude of this effect for non-compounds from that of compounds, separately for each perceptual domain. We used the same Bayesian procedure described above, this time to compare the hypothesis that the difference between compounds and non-compounds in the magnitude of the old/rearranged effect does not differ for within-domain and cross-domain word-pairs, with the hypothesis that the old/rearranged effect for compounds versus non-compounds is greater for within-domain than cross-domain word-pairs. This analysis also supported the null hypothesis, which was preferred by a Bayes factor of 5.26, thus providing moderate evidence against our third prediction.

#### 3.2.2 ERP results: Comprehensive analyses

Topographic maps of the old/rearranged effect in the various conditions are shown in
Figure 2, panels A-B. Figure 2 panels C-D shows grand-mean ERPs for each condition, in a representative fontal electrode (Fz; where associative recognition effects in the early time window were maximal) and a representative left-central electrode (C3; where the effects in the late time window were maximal). As can be seen, in the early time window, differences related to associative recognition (i.e., less negativity for intact versus rearranged pairs) emerged at frontal locations for compound words, but were absent for non-compounds, and were more pronounced for cross-domain than for within-domain word-pairs. In the late time window, differences related to associative recognition were maximal at the left-central location, and emerged for both compounds and non-compounds (though were more pronounced for compounds), and for both within-domain and cross-domain pairs (though were more pronounced for cross-domain pairs).

#### 3.2.2.1 Early time window (300-500ms)

In the early time window, the analysis revealed a marginally significant main effect of response type *F*(1, 16) = 3.2, *p* = .09, 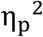 = .17, a marginally significant interaction between word-pair type and response type, *F*(1, 16) = 3.7, *p* = .07, 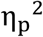 = .19, stemming from a significant old/rearranged effect (greater negativity for rearranged than old responses) for compounds, *t*(16) = 3.03, *p* = .008, but not for non-compounds, *t*(16) = -0.21, *p* = .835.

The analysis further revealed significant interactions between perceptual domain, response type, and laterality, *F*(2, 32) = 7.44, *p* = .002, 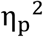 = 0.32, and between perceptual domain, response type, anteriority, and laterality, *F*(4, 64) = 2.67, *p* = .04, 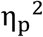 = 0.14. Follow-up analysis of the latter interaction revealed that for within-domain word-pairs, the old/rearranged effect was marginal in frontal sites (F3, *p* = .058; Fz, *p* = .067; F4, *p* = .078), and absent in other locations. In contrast, for cross-domain word-pairs, the old/rearranged effect was significant (or marginally significant) in right-frontal and right-central locations (Fz, *p* = .006; F4 = .075; Cz, *p* = 0.015; C4, *p* = 0.004).

Finally, the analysis revealed an interaction between response type and laterality, *F*(2, 32) = 4.34, *p* = .021, 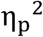 = 0.21, stemming from a significant old/rearranged effect in midline locations, *t*(16) = 2.39, a marginal effect in right-lateralized locations, *t*(16) = 1.93, *p* = .07, but no effect in left-lateralized locations, *t*(16) = 0.94, *p* = .361, and a significant interaction between response type and anteriority, *F*(2, 32) = 15.93, *p* < 0.0001, 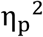 = 0.50, stemming from a significant old/rearranged effect in frontal and central locations, *t*(16) = 3.01, *p* = .008; *t*(16) = 2.37, *p* = .031, but no effect in posterior locations, *t*(16) = -0.59, *p* = .564.

#### 3.2.2.2 Late time window (500–800 ms)

The analysis of old and rearranged responses in the late time window revealed a significant main effect of response type *F*(1, 16) = 34.89, *p* < .0001, 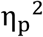= 0.69, and significant interactions between response type and anteriority, *F*(2,32) = 10.93, *p* < .0001, 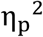= 0.41, and between response type and laterality, *F*(2, 32) = 4.73, *p* = .016, 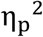= 0.23, indicating that although the old/rearranged effect was widespread and significant in all locations (*ps* < 0.0001), it was maximal in left and central locations.

Furthermore, the analysis revealed significant interactions between word-pair type and response type, *F*(1, 16) = 10.56, *p* = .005, 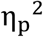= 0.40, and between word-pair type, response type, and anteriority, *F*(2, 32) = 4.96, *p* = .013, 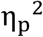= 0.24. Follow-up analysis of the latter interaction revealed that in posterior locations, the old/rearranged effect was significant for compounds (*p* < .0001), but not for non-compounds (*p* > .05), whereas in frontal and central locations the old/rearranged effect emerged for both compounds (*p* < 0.0001, *p* < 0.0001) and non-compounds (*p* = 0.011, *p* = 0.042), but was greater for the former.

Finally, significant interactions were revealed between perceptual domain, response type, and laterality, *F*(2, 32) = 5.89, *p* = .007, 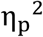= 0.27, and between perceptual domain, response type, anteriority, and laterality, *F*(4, 64) = 3.19, *p* = .019, 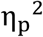= 0.17. Follow-up analysis of the 4-way interaction revealed that while for cross-domain pairs the old/rearranged effect was significant in all locations (*ps* < 0.05), for within-domain pairs, the effect was absent in the right-posterior location *(p* = .415), and was significant, but attenuated, in other locations (*ps* < .05).

## 4 Discussion

In the current study, an associative recognition memory task was employed to explore whether episodic associations between compounds versus non-compounds and between within-domain versus cross-domain word-pairs differentially recruit familiarity- and recollection-based recognition, as indexed by their putative electrophysiological signatures. The critical hypothesis underlying our current study was that the ability of associations to be processed in a unitized fashion may promote the contribution of familiarity to their associative recognition. Therefore, we focused on the early frontal old/rearranged effect, conventionally interpreted as the putative ERP correlate of associative familiarity, but also evaluated other associative recognition effects in the early and late time windows. Our data provide novel evidence for a multiplicity of processes supporting associative recognition, and afford new insights regarding two theoretical frameworks that account for mnemonic unitization effects—the DD view and the LOU account.

Our first key finding was that associative recognition of compounds (but not of non-compounds) was associated with modulation of an early frontal effect—the putative electrophysiological correlate of familiarity—as indicated by decreased frontal negativity for old compared to rearranged compounds in the early time window. Several previous electrophysiological studies (Bader et al., 2010; Diana et al, 2011; Guillaume & Etienne, 2015; Jäger et al., 2006; Jäger et al., 2010; Kamp et al., 2016; Kriukova et al., 2013; Li, Mao, Wang, & Guo, 2017; Lyu, Wang, Mao, Li, Guo, 2018; Rhodes & Donaldson, 2008; Tibon et al., 2014a; Tibon et al., 2014b; Zheng et al., 2015) support the notion that familiarity is enabled for unitized associations. However, only two prior electrophysiological investigations of unitization effects at the higher end of the LOU continuum (where stimuli already share strong associative relationships prior to the experiment) employed a design that allows conclusive attribution of mnemonic effects to associative recognition (Kriukova et al., 2013, Zheng et al., 2015). Other studies do not afford direct examination of associative effects, either because the experimental conditions in the design were not fully balanced (e.g., related stimuli were rearranged into unrelated retrieval pairs, Bader et al., 2010; Li et al., 2017; Rhodes & Donaldson, 2008; Tibon et al., 2014b; Wang et al., 2016), or because the analyses contrasted old/new responses, rather than old/rearranged responses (Bader et al. 2010; Rhodes & Donaldson, 2008; Wiegand, Bader, & Mecklinger, 2010). Notably, in such cases, the early frontal negativity can be interpreted in terms of semantic congruency rather than episodic memory (e.g., see discussion in Tibon, Cooper, & Greve, 2017), or in terms of item rather than associative recognition. Crucially, in the current study, old compounds were contrasted with rearranged compounds, and old non-compounds were contrasted with rearranged non-compounds. Therefore, any retrieval effects can be attributed to the influence of a pre-existing association on the *episodic* memory of the pairs (that is, whether the pair presented at test was encoded before, during the study phase). With this, our study joins a handful of previous electrophysiological studies (Kriukova et al., 2013, Zheng et al., 2015), by conclusively showing that retrieval episodic associations between stimuli that share strong associative relationships prior to the experiment, is associated with an early ERP component strongly linked to familiarity.

Our second key finding was that, in accord with the prediction of the LOU account and contrary to the prediction of the DD view, modulation of the early frontal effect by old versus rearranged pairs was not greater for within-domain relative to cross-domain associations. This was further evident in our Bayesian analysis which preferred the null hypothesis of no difference. Our finding that within-domain and cross-domain associations equally modulate the early frontal effect (thereby suggesting that associative familiarity is available for both types of perceptual domains), thus agrees with the proposal of the LOU account, that the unitization process is capable of operating at a fairly abstract level, and is therefore also available across different stimulus modalities or domains. Not only was there no unitization advantage for within-domain word-pairs, in the current study we observed a somewhat reversed pattern: our behavioral data revealed that accuracy (measured by associative Pr, see Table 2) was marginally better for cross-domain than for within-domain word-pairs. This was accompanied by the results of our comprehensive ERP analysis, showing a greater frontal associative recognition effect for cross-domain than within-domain word-pairs.

This reversed pattern contrasts with some previous studies, showing greater unitization advantage / familiarity-based retrieval / non-hippocampal based retrieval for within-domain associations (Bastin et al., 2010; Borders et al., 2017; Mayes et al., 2014, Tibon et al., 2014a; Troyer et al., 2011; Vergha-Khadem et el., 1997), but agrees with other studies, showing that cross-domain associations can also benefit from unitization (Harlow et al., 2010; Park & Rugg, 2011; Parks & Yonelinas, 2015; Tzurizziani et al., 2004). In the current study, we tested the hypothesis (formulated as our third prediction) that this discrepancy arises because the relation between stimuli presentation (within/cross-domain) and unitization effects is mediated by preexisting associative relations between the stimuli, such that only stimuli that already bear a strong association (such as compound words) would benefit from a within-domain presentation. However, our study did not confirm this hypothesis, and did not yield the predicted interaction between word type and perceptual domain on the magnitude of the early frontal old/rearranged effect. An alternative hypothesis to the one we set to test in the current study, is that relatively simple stimuli such as words are more easily unitized within-domain compared to relatively complex stimuli such as fractals (Parks & Yonelinas, 2015). However, our current study does not support this hypothesis either, since the stimuli in our case were word-pairs which are considered by Parks & Yonelinas (2015) to be relatively simple. It is therefore yet to be determined why some studies, but not in others, show greater unitization effects for within-domain associations.

In addition to these key findings, emerging from our pre-defined predictions, our comprehensive data analysis further revealed an associative recognition effect in the late time window. Importantly, in this time window, the effect was maximal in central-posterior locations (adhering to the typical distribution of the putative electrophysiological correlate of recollection), and was observed for all conditions. This indicates that unlike the early frontal effect, which is only available for unitized associations (in our case – compound words), the late posterior effect is associated with associative recognition, regardless the nature of the association—as would be expected from an electrophysiological correlate of recollection. Nevertheless, the magnitude of the late effect differed across the various conditions. Firstly, the effect was greater for compounds versus non-compounds. This pattern is in accord with previous studies, which generally agree that congruent stimuli combinations, such as word compounds (vs. non-compounds), support high Levels of Processing (LOP; Craik & Lockhart, 1972), yielding rich and elaborate encoding. In turn, during retrieval, this elaborated encoding supports high levels of recollection (summarized by Craik, 2002), which in our case, was indicated by greater late posterior positivity for old than rearranged pairs. Secondly, the late posterior effect was greater for cross-domain than within-domain associations. Given previously reported links between the late posterior component and mnemonic related activation in the posterior parietal lobe (PPC; Vilberg & Rugg, 2009), this pattern agrees with the suggestion linking PPC activation with increased recollection due to multimodal integration of retrieved stimuli (Ben-Zvi, Soroker, & Levy, 2015; Bonnici, Richter, Yazar, & Simons, 2016; Shimamura, 2011, Yazar et al., 2017, but see Tibon et al., 2014a; Tibon & Levy, 2014 for a different pattern of results). However, given the poor spatial resolution of EEG, and the fact that the current study did not localize the source of the mnemonic ERP effects, any conclusions regarding neural sources of these effects should be treated with caution.

In our study, the distinction between the two processes supporting recognition memory—familiarity and recollection—is built of the temporal and spatial distribution of the early and late old/rearranged ERP effects. Like many other studies (e.g., Bader et al., 2010; Guillaume & Etienne, 2015; Jäger et al., 2006; Kamp et al., 2016; Rhodes & Donaldson, 2008; Tibon et al., 2014b; Zheng et al., 2015, to list just a few), we link the early frontal negativity with familiarity, and the late posterior positivity with recollection. While this type of reverse inference has its limitations (Poldrack, 2006), our interpretations build on decades of intensive research that strongly associates these ERP components with the particular memory processes reported here (reviewed by Mecklinger, 2000, Rugg & Curran, 2007, Wilding & Ranganath, 2011). In any event, however, even if the links made in our study between electrophysiological measures and specific recognition processes are not entirely conclusive (though strongly suggestive), our data clearly point to a neural distinction, whereby the contribution of the early ERP effect to associative recognition is limited to unitized associations, but the contribution of the late ERP effect is available for all types of associative information.

In summary, examining the electrophysiological correlates of episodic associative recognition, we report evidence suggesting that recognition of compounds amenable to unitization can be differentially supported by associative familiarity, regardless of whether both words comprising the compound were presented to the same sensory modality, or to different sensory modalities. These results reinforce the importance of pre-existing associative relationships between episodically associated memoranda, in determining how their cooccurrence is experienced and subsequently remembered. Furthermore, our finding that unitization can support cross-domain associations suggests that it can act as a useful encoding strategy across a broad range of experimental conditions and materials.

## Acknowledgements

The present study was supported by the National Natural Science Foundation of China (31671127), Support by "Capacity Building for Sci-Tech Innovation – Fundamental Scientific Research Funds (No. 025185305000/200). RT is supported by a British Academy Postdoctoral Fellowship (grant SUAI/028 RG94188).

The authors declare no conflict of interest.

